# The effect of walking with reduced trunk motion on dynamic stability in healthy adults

**DOI:** 10.1101/2022.11.03.515025

**Authors:** Tom J.W. Buurke, Lotte van de Venis, Noël Keijsers, Jorik Nonnekes

**Affiliations:** University of Groningen, University Medical Center Groningen, Department of Human Movement Sciences, Groningen, The Netherlands; KU Leuven, Department of Movement Sciences, Leuven, Belgium; Radboud University Medical Centre, Nijmegen, The Netherlands; Donders Institute for Brain, Cognition and Behaviour; Center of Expertise for Parkinson & Movement Disorders; Department of Rehabilitation; Sint Maartenskliniek, Department of Research, Nijmegen, The Netherlands; Radboud University, Nijmegen, The Netherlands; Donders Institute for Brain, Cognition and Behaviour; Department of Sensorimotor Neuroscience

**Keywords:** Locomotion, Gait stability, Trunk rigidity, Margin of stability, Parkinson’s Disease

## Abstract

**Background:** Most people with Parkinson’s Disease (PD) walk with a smaller mediolateral base of support (BoS) compared to healthy people, but the underlying mechanisms remain unknown. According to the extrapolated center of mass (XCoM) concept, a decrease in mediolateral XCoM excursion would require a smaller mediolateral BoS to maintain a constant margin of stability (MoS) and remain stable. As people with PD typically walk with reduced trunk motion, we hypothesized that the mediolateral MoS might stay the same despite a smaller BoS.

**Research question:** As proof of principle, we assess whether walking with reduced trunk motion results in a smaller step width in healthy adults, without altering the mediolateral MoS.

**Methods:** Fifteen healthy adults walked on a treadmill at preferred comfortable walking speed in two conditions. First, the ‘regular walking’ condition without any instructions, and second, the ‘reduced trunk motion’ condition with the instruction: ‘Keep your trunk as still as possible’. Treadmill speed was kept the same in the two conditions. Trunk kinematics, step width, mediolateral XCoM excursion and mediolateral MoS were calculated and compared between the two conditions.

**Results:** Walking with the instruction to keep the trunk still significantly reduced trunk kinematics. Walking with reduced trunk motion resulted in significant decreases in step width and mediolateral XCoM excursion, but not in the mediolateral MoS. Furthermore, step width and mediolateral XCoM excursion were strongly correlated during both conditions (r=0.887 and r=0.934).

**Significance:** This study shows that walking with reduced trunk motion leads to a gait pattern with a smaller BoS in healthy adults, without altering the mediolateral MoS. Our findings indicate a strong coupling between CoM motion state and the mediolateral BoS. We expect that people with PD who walk narrow-based, have a similar mediolateral MoS as healthy people, which will be further investigated.

## Introduction

People with a neurological gait disorder (e.g. ataxia due to polyneuropathy) typically walk with an increased mediolateral base of support (BoS) compared to healthy people [1,2]. However, there are neurological patient groups in which a smaller BoS is seen, including people with spastic paraplegia and Parkinson’s disease (PD) [3]. PD is characterized by motor and non-motor symptoms, of which rigidity, bradykinesia, and tremor are the key motor features [4]. A smaller mediolateral BoS in people with spastic paraplegia is related to hip adductor spasticity [5], but the mechanisms underlying narrow-based gait in people with PD remain to be explored.

The size of the mediolateral BoS during walking is related to the mediolateral center of mass (CoM) position and velocity [6–8], captured in the extrapolated center of mass (XCoM) concept [6,9]. The XCoM describes the CoM motion state, based on CoM position and CoM velocity normalized to a constant factor related to posture [6,10]. This concept states that an increase in mediolateral XCoM position requires a proportionate increase in the mediolateral BoS to maintain a similar distance between the mediolateral XCoM and mediolateral BoS, i.e., the Margin of Stability (MoS), in order to remain stable. In PD, axial rigidity may decrease mediolateral CoM motion during walking. Therefore, one could hypothesize that narrow-based gait in people with PD may be related to trunk rigidity, as walking with reduced trunk motion could decrease mediolateral CoM motion and therefore require a smaller step width to maintain a constant MoS and remain stable [9].

Previously, Arvin and colleagues showed that constrained trunk motion during walking by a corset resulted in decreases in mediolateral CoM position, mediolateral CoM velocity and step width [11]. However, the use of an external device to constrain trunk motion may limit the ecological validity. Here, we assessed whether walking with reduced trunk motion results in a narrow-based gait pattern in healthy adults. Therefore, we instructed healthy participants to reduce trunk motion while walking on a treadmill. We hypothesized that walking with reduced trunk motion leads to reduced mediolateral XCoM excursion coinciding with a smaller in step width, which in turn leads to an unaltered mediolateral MoS compared to regular walking. Furthermore, we assessed the correlational relationship between mediolateral XCoM excursion and step width, to gain further insight into the mechanisms underlying narrow-based gait. Based on literature [11], we hypothesized a relationship between mediolateral XCoM excursion and step width. In addition, we expect a relationship between the decrease in mediolateral XCoM excursion and decrease in step width due to reduced trunk motion.

## Methods

### Participants

Fifteen healthy adults (5 female, mean ± SD age: 44 ± 13 years, height: 1.79 ± 0.08 m, weight: 78 ± 11 kg) volunteered to participate in this study. Participants were included if they were between 18 and 70 years and did not have any reported pathologies that are known to impact balance or gait. Participants provided written informed consent before study onset and the procedures of this study were in line with the Declaration of Helsinki [12] and local ethical regulations.

### Experimental protocol

Participants walked on an instrumented dual-belt treadmill (Motek, Amsterdam, NL), during which they were not allowed to hold on to the treadmill’s handrails to prevent the effects of external stabilization on gait [13]. To provide safety, participants were fitted with a harness attached to the ceiling, which did not constrain movement or provide body weight support. All participants walked at their preferred comfortable walking speed (mean ± SD: 1.30 ± 0.11 m s^−1^). To find the preferred comfortable walking speed, belt speed was manually increased until the participant indicated it was comfortable. Belt speed was then increased with 0.3 m s^−1^ and slowly decreased until the participant again indicated it was comfortable. The average of both speeds was used to set the preferred comfortable walking speed of each participant [14]. The treadmill maintained a constant speed, which was kept the same in two conditions. First, the ‘regular walking’ condition without any instructions, and second, the ‘reduced trunk motion’ condition with the instruction: ‘Keep your trunk as still as possible’. The instruction was continuously repeated by the experimenter during the reduced trunk motion condition. The order of the two conditions was the same for all participants. Both conditions lasted three minutes and the final two minutes of each condition were selected for further analysis to exclude task familiarization effects in the first minute of each new condition.

### Data acquisition

A 10-camera motion capture system (Vicon Motion Systems Ltd., Oxford, UK) recorded 3D kinematics at 100 Hz. Markers were placed according to the standard Plug-in Gait model for the upper and lower body [15]. Subsequently, the kinematic data were processed in Vicon Nexus 2.10.1 (Vicon Motion Systems Ltd., Oxford, UK). The treadmill’s embedded force plates captured 6D ground reaction forces (N, Nm) and 2D center of pressure positions (m) at 2000 Hz.

### Data analysis

The data were analyzed in MATLAB r2020b (MathWorks Inc., Natick, MA, USA). Gait events were detected by finding the local peaks and valleys in the anteroposterior direction of foot marker data [16]. Trunk obliquity (deg) and trunk rotation (deg) were extracted from the Vicon Plug-in Gait model and defined in absolute values [15]. The maximum range of motion in trunk obliquity and trunk rotation was calculated for each stride to define trunk kinematics. Ground reaction forces and center of pressure positions were low-pass filtered at 15 Hz with a 2^nd^ order Butterworth filter. Step width (m) was calculated as the difference in mediolateral center of pressure position between ipsilateral and contralateral toe-off for each step, to match the definition of the mediolateral BoS in the calculation of the mediolateral MoS below. The mediolateral CoM position (m) was calculated by combining the twice-integrated and high-pass filtered (0.2 Hz) mediolateral GRF divided by the participant’s weight with the low-pass filtered (0.2 Hz) mediolateral center of pressure [17–19]. The mediolateral XCoM (m) was calculated according to Equation 1, where l is pendulum length (m), defined as leg length multiplied by 1.2 [6], g is the gravitational acceleration (9.81 m s^−2^) and vCoM is CoM velocity (m s^−1^) [6]. The mediolateral XCoM excursion was defined as the difference in mediolateral XCoM position between the ipsilateral and contralateral toe-off for each stride. The mediolateral MoS (m) was calculated as the distance between the mediolateral CoP position and the mediolateral XCoM position at the time instance of contralateral toe-off (*j)* for each step (*i)* [6], as in Equation 2.

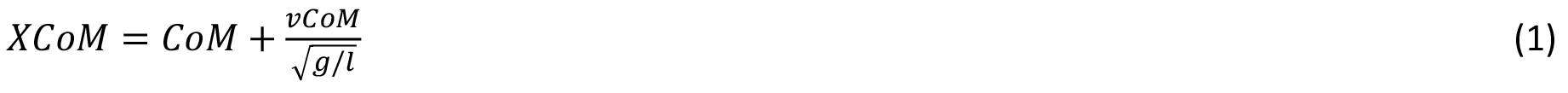

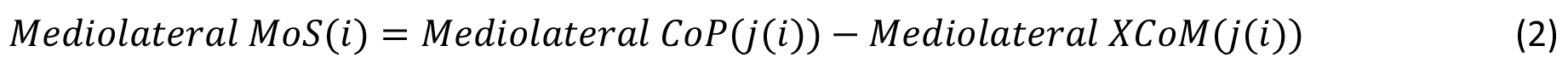

### Statistical analysis

To assess the effect of walking with reduced trunk motion on trunk kinematics and gait stability parameters, we conducted multiple statistical tests in MATLAB r2020b with the Statistics and Machine Learning Toolbox (MathWorks Inc., Natick, MA, USA). The average values of trunk obliquity, trunk rotation, step width, mediolateral XCoM excursion, and mediolateral MoS were calculated over all strides or steps for each participant. The data were normally distributed. Statistical significance was set at an alpha of 0.05 with a Bonferroni correction to adjust for multiple comparisons, leading to a corrected alpha of 0.00625 [20].

To establish whether the instruction to walk with reduced trunk motion led to decreased trunk kinematics during walking, we performed paired t-tests to compare trunk obliquity and trunk rotation between the regular walking condition and the reduced trunk motion condition. Then, to assess the effect of walking with reduced trunk motion on gait stability parameters, we performed paired t-tests to compare step width, mediolateral XCoM excursion, and the mediolateral MoS between the two conditions. To gain insight into the relationship between mediolateral XCoM excursion and step width, we calculated Pearson’s correlation coefficient between mediolateral XCoM excursion and step width during the regular walking condition and the reduced trunk motion condition. Finally, we assessed whether changes in mediolateral XCoM excursion due to reduced trunk motion were related to changes in step width due to reduced trunk motion. To this end, we calculated Pearson’s correlation coefficient between the difference in mediolateral XCoM excursion and the difference in step width between the regular walking condition and the reduced trunk motion condition.

## Results

The data of a representative subject illustrates that walking with reduced trunk motion decreased trunk kinematics (Figure 1A). At group level, walking with the instruction to ‘keep your trunk as still as possible’ significantly decreased trunk obliquity (Figure 2A, mean ± SD regular walking: 3.14 ± 1.17 deg, reduced trunk motion: 1.49 ± 0.37 deg, t = 6.151, p < 0.001) and trunk rotation (Figure 2B, mean ± SD regular walking: 6.43 ± 2.40 deg, reduced trunk motion: 4.44 ± 1.25 deg, t = 3.787, p = 0.002) compared to regular walking without any instructions.

**Figure 1.**
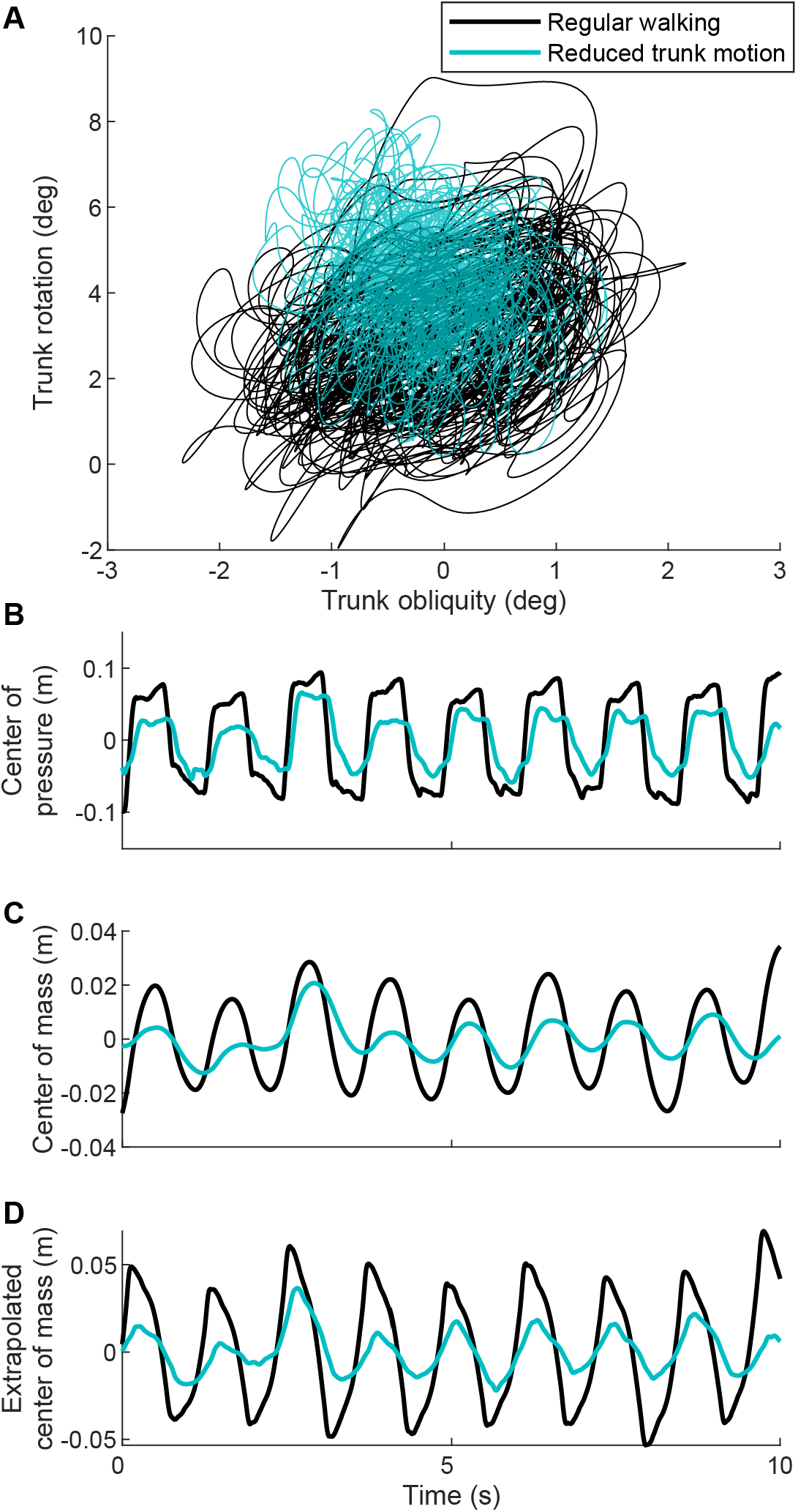
Data of a single participant that shows typical behavior during the regular walking condition (black) and the reduced trunk motion condition (cyan). A) Trunk rotation vs trunk obliquity for the duration of the whole experiment. B) Mediolateral center of pressure position, C) mediolateral center of mass position and D) mediolateral extrapolated center of mass position during ten seconds of the experiment.

**Figure 2.**
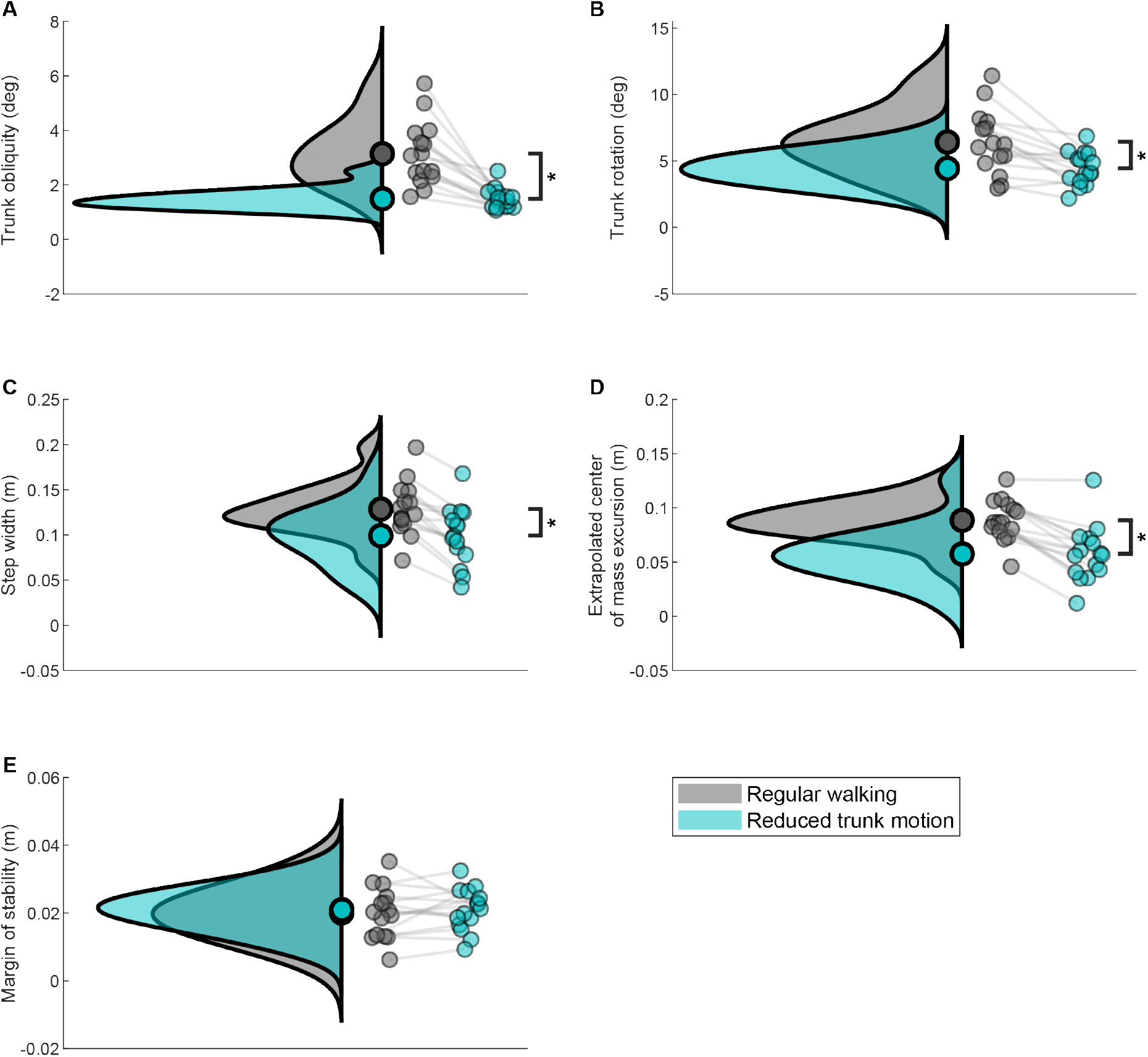
Vertical raincloud plots [21] of the group data during the regular walking condition (grey) and the reduced trunk motion condition (cyan). Subplots show A) trunk obliquity, B) trunk rotation, C) step width, D) mediolateral extrapolated center of mass excursion and E) mediolateral margin of stability. The left half of each raincloud plot shows the group distribution and group mean for each condition. The right half of each raincloud plot shows the data for individual participants connected by a line between the two conditions. Asterisks indicate significant within-group differences between the conditions (p < 0.05).

Walking with reduced trunk motion decreased the mediolateral BoS, mediolateral CoM excursion and mediolateral XCoM excursion in a representative subject (Figure 1B,C,D). At group level, walking with reduced trunk motion decreased step width (Figure 2C, mean ± SD regular walking: 0.129 ± 0.029 m, reduced trunk motion: 0.099 ± 0.033 m, t = 6.867, p < 0.001) and mediolateral XCoM excursion (Figure 2D, mean ± SD regular walking: 0.089 ± 0.019 m, reduced trunk motion: 0.058 ± 0.026 m, t = 7.325, p < 0.001). Interestingly, there was no difference in the mediolateral MoS between the two conditions (Figure 2E, mean ± SD regular walking: 0.020 ± 0.008 m, reduced trunk motion: 0.021 ± 0.006 m, p = 0.563).

Pearson’s correlation analyses showed a significant positive relationship between mediolateral XCoM excursion and step width in the regular walking (r = 0.887, p < 0.001) and the reduced trunk motion condition (r = 0.934, p < 0.001, Figure 3A). Furthermore, we found a significant positive correlation between the decrease in step width and the decrease in mediolateral XCoM excursion due to reduced trunk motion (r = 0.731, p = 0.002, Figure 3B).

**Figure 3.**
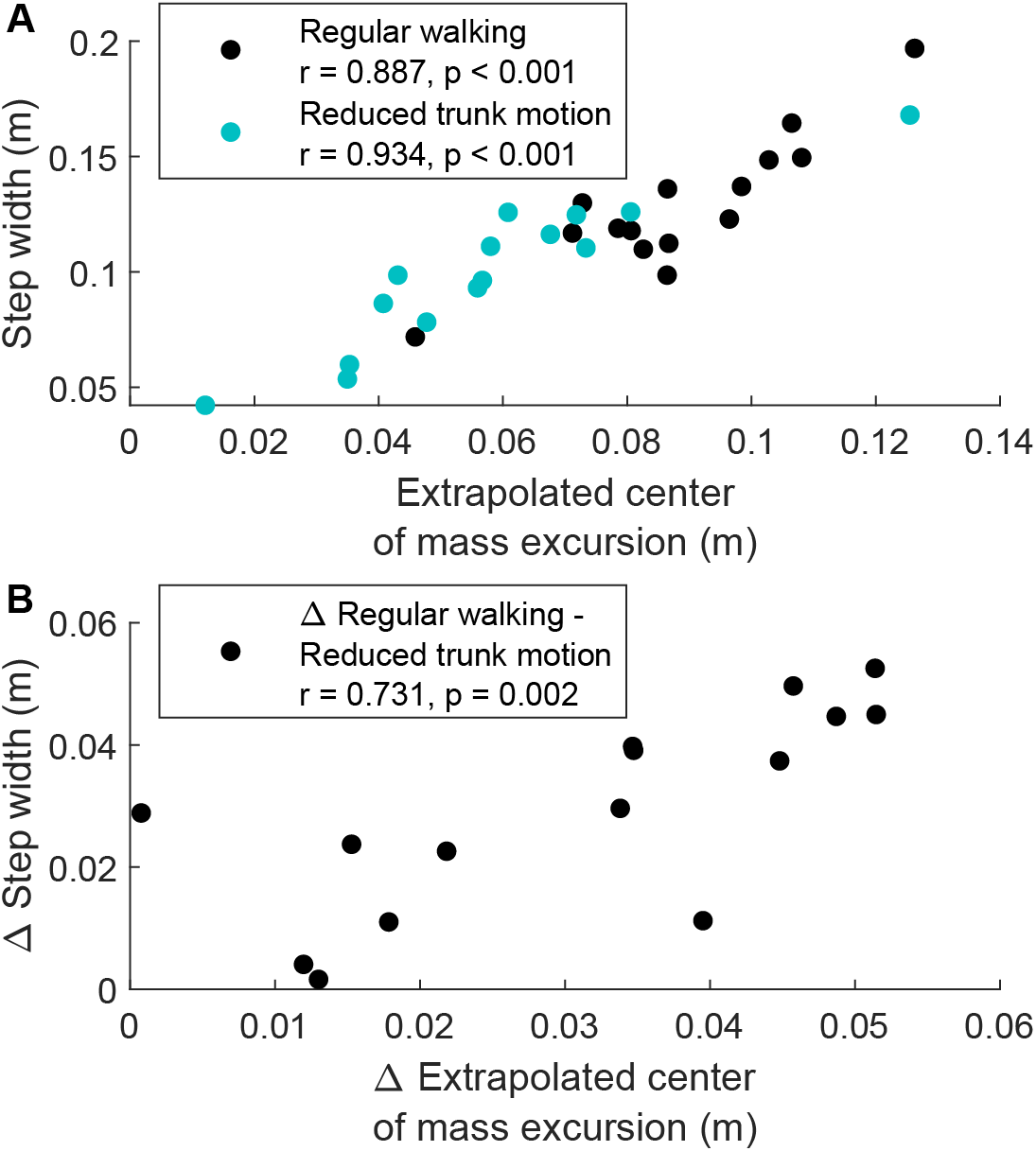
Scatter plots of the relationship between extrapolated mediolateral center of mass excursion and step width. Subplots show A) the regular walking condition (black) and the reduced trunk motion condition (cyan), and B) the change in extrapolated center of mass excursion and step width due to reduced trunk motion. Each data point represents an individual participant.

## Discussion

In this study, healthy adults successfully decreased their trunk kinematics when walking with the instruction to reduce trunk motion. Walking with reduced trunk motion resulted in decreases in step width and mediolateral XCoM excursion. As hypothesized, we found no changes in the mediolateral MoS during walking with reduced trunk motion compared to regular walking. Furthermore, we found a strong relationship between mediolateral XCoM excursion and step width during regular walking and during walking with reduced trunk motion, as well as in the decrease in mediolateral XCoM excursion and step width due to walking with reduced trunk motion. These results show that walking with reduced trunk motion leads to a narrow-based gait pattern in healthy adults, without altering the mediolateral MoS. Furthermore, our findings indicate a strong coupling between CoM motion state and the mediolateral BoS [9].

Previously, it has shown that walking with smaller step width by walking on a projected line on the treadmill decreases mediolateral CoM kinematics [22]. Furthermore, decreased mediolateral CoM motion through external stabilization resulted in a decrease in step width [23–25], which can be explained by a strong relationship between the CoM motion state and foot placement [26]. Finally, walking with a corset that constrained trunk motion resulted in smaller mediolateral CoM position, CoM velocity and step width [11]. Together with our results, these findings emphasize the strong coupling between CoM motion state and the mediolateral BoS during walking.

The exact mechanisms underlying the coupling between the CoM motion state and mediolateral BoS remain to be unraveled. Possibly, the coupling reflects a ‘simple constant offset control rule for stable walking’ [9]. This is supported by the finding that mediolateral foot placement adjustments after an external perturbation applied to the pelvis were found to be proportionate to the mediolateral CoM velocity [27], where automatic hip abductor responses may be partially responsible for these adjustments [28]. In support of this hypothesis, it was found that over 80% of lateral foot placement variance can be predicted by CoM motion state at mid-stance and 89% at terminal swing [26]. Furthermore, it was shown that a CoM feedback model including CoM position, velocity and acceleration can explain muscle responses during postural balance responses, which suggest that the central nervous system uses a multisensory estimate of CoM motion state to control balance [29].

These studies leave the question how strong the coupling between the CoM motion state and the mediolateral BoS is when task and environmental demands change. For instance, humans are known to increase the aforementioned constant offset, i.e. the mediolateral MoS, in anticipation of gait perturbations [30] by changing stepping parameters such a step width and stance time [10,31], potentially at the cost of metabolic efficiency [32]. It is unclear exactly how the trade-off between gait stability and metabolic efficiency is evaluated in human walking, e.g., when uncertainty about the CoM motion state is augmented. In future research we aim to gain further insight into these underlying mechanisms of dynamic stability in human walking.

It remains the question whether walking with the instruction to reduce trunk motion fully resembles trunk rigidity in people with PD. Reduced trunk motion during walking has been documented in PD using a variety of outcomes [33–36]. While outcome parameters and absolute values differ between these studies and ours, the decreases in trunk obliquity and trunk rotation due to reduced trunk motion in the current study are comparable to the differences between healthy adults and people with PD in trunk obliquity [33,34,36] and trunk rotation [35,36]. As we aimed to test a proof of principle, we believe that the choice for an instruction aimed at trunk motion was sensible and valid. In the future, we aim to assess whether trunk rigidity indeed underlies narrow-based gait in people with PD. Recently, it was shown that the sensorimotor utilization of CoM feedback during standing [29] is intact in people with PD, but may be disinhibited in antagonistic muscles during perturbations [37]. This emphasizes that further studies into the coupling between mediolateral CoM motion state and the mediolateral BoS during walking in people with PD are necessary to understand the underlying mechanisms of narrow-based gait.

Here, we aimed to test a proof of principle for the underlying mechanisms of narrow-based gait in people with PD. We mimicked Parkinsonian trunk rigidity by asking healthy participants to keep their trunk as still as possible while walking on a treadmill. This study shows that the proof of principle holds true; narrow-based gait is the result of reduced trunk motion to maintain a similar mediolateral MoS. In future research we will investigate whether this principle holds in people with PD and whether uncertainty about the CoM motion state increases the mediolateral MoS during walking.

## CRediT authorship contribution statement

**Tom J**.**W. Buurke**: Conceptualization, Methodology, Software, Formal analysis, Data curation, Writing – original draft, Visualization. **Lotte van de Venis**: Methodology, Investigation, Data curation, Writing – review & editing. **Noël Keijsers**: Methodology, Investigation, Writing – review & editing, Supervision. **Jorik Nonnekes**: Conceptualization, Methodology, Investigation, Writing – review & editing, Supervision, Project administration.

## References

[1] G. Chen, C. Patten, D.H. Kothari, F.E. Zajac, Gait differences between individuals with post-stroke hemiparesis and non-disabled controls at matched speeds, Gait Posture. 22 (2005) 51–56. https://doi.org/10.1016/j.gaitpost.2004.06.009.

[2] S. Chakraborty, A. Nandy, T.M. Kesar, Gait deficits and dynamic stability in children and adolescents with cerebral palsy: A systematic review and meta-analysis, Clinical Biomechanics. 71 (2020) 11–23. https://doi.org/10.1016/j.clinbiomech.2019.09.005.

[3] J. Nonnekes, R.J.M. Goselink, E. Růžička, A. Fasano, J.G. Nutt, B.R. Bloem, Neurological disorders of gait, balance and posture: a sign-based approach, Nat Rev Neurol. 14 (2018) 183–189. https://doi.org/10.1038/nrneurol.2017.178.

[4] B.R. Bloem, M.S. Okun, C. Klein, Parkinson’s disease, The Lancet. 397 (2021) 2284–2303. https://doi.org/10.1016/S0140-6736(21)00218-X.

[5] J. Nonnekes, B. van Lith, B.P. van de Warrenburg, V. Weerdesteyn, A.C.H. Geurts, Pathophysiology, diagnostic work-up and management of balance impairments and falls in patients with hereditary spastic paraplegia, J Rehabil Med. 49 (2017) 369–377. https://doi.org/10.2340/16501977-2227.

[6] A.L. Hof, M.G.J. Gazendam, W.E. Sinke, The condition for dynamic stability, J. Biomech. 38 (2005) 1–8. https://doi.org/10.1016/j.jbiomech.2004.03.025.

[7] Y.C. Pai, J. Patton, Center of mass velocity-position predictions for balance control, J. Biomech. 30 (1997) 347–354. https://doi.org/10.1016/S0021-9290(96)00165-0.

[8] M.A. Townsend, Biped gait stabilization via foot placement, Journal of Biomechanics. 18 (1985) 21–38. https://doi.org/10.1016/0021-9290(85)90042-9.

[9] A. Hof L., The ‘extrapolated center of mass’ concept suggests a simple control of balance in walking, Hum. Mov. Sci. 27 (2008) 112–125. https://doi.org/10.1016/j.humov.2007.08.003.

[10] L. Hak, H. Houdijk, F. Steenbrink, A. Mert, P. van der Wurff, P.J. Beek, J.H. van Dieën, Speeding up or slowing down?: Gait adaptations to preserve gait stability in response to balance perturbations, Gait Posture. 36 (2012) 260–264. https://doi.org/10.1016/j.gaitpost.2012.03.005.

[11] M. Arvin, J.H. van Dieën, S.M. Bruijn, Effects of constrained trunk movement on frontal plane gait kinematics, Journal of Biomechanics. 49 (2016) 3085–3089. https://doi.org/10.1016/j.jbiomech.2016.07.015.

[12] World Medical Association, World Medical Association declaration of Helsinki: ethical principles for medical research involving human subjects, JAMA. 310 (2013) 2191–2194. https://doi.org/10.1001/jama.2013.281053.

[13] T.J.W. Buurke, C.J.C. Lamoth, L.H.V. van der Woude, R. den Otter, Handrail Holding during Treadmill Walking Reduces Locomotor Learning in Able-Bodied Persons, IEEE Trans. Neural Syst. Rehabil. Eng. 27 (2019) 1753–1759. https://doi.org/10.1109/TNSRE.2019.2935242.

[14] L. van de Venis, B.P.C. van de Warrenburg, V. Weerdesteyn, B.J.H. van Lith, A.C.H. Geurts, J. Nonnekes, Improving gait adaptability in patients with hereditary spastic paraplegia (Move-HSP): study protocol for a randomized controlled trial, Trials. 22 (2021) 32. https://doi.org/10.1186/s13063-020-04932-9.

[15] V.M.S. Limited, Plug-in Gait Reference Guide, (2021). https://docs.vicon.com/display/Nexus212/PDF+downloads+for+Vicon+Nexus?preview=/133828966/133829807/Plug-in%20Gait%20Reference%20Guide.pdf (accessed January 19, 2022).

[16] J.A. Zeni, J.G. Richards, J.S. Higginson, Two simple methods for determining gait events during treadmill and overground walking using kinematic data, Gait & Posture. 27 (2008) 710–714. https://doi.org/10.1016/j.gaitpost.2007.07.007.

[17] T.J.W. Buurke, C.J.C. Lamoth, D. Vervoort, L.H.V. van der Woude, R. den Otter, Adaptive control of dynamic balance in human gait on a split-belt treadmill, J. Exp. Biol. 221 (2018) jeb174896. https://doi.org/10.1242/jeb.174896.

[18] A.L. Hof, Comparison of three methods to estimate the center of mass during balance assessment, J. Biomech. 38 (2005) 2134–2135. https://doi.org/10.1016/j.jbiomech.2005.03.029.

[19] H.M. Schepers, E.H.F. van Asseldonk, J.H. Buurke, P.H. Veltink, Ambulatory estimation of center of mass displacement during walking, IEEE Trans. Biomed. Eng. 56 (2009) 1189–1195. https://doi.org/10.1109/TBME.2008.2011059.

[20] C. Bonferroni, Teoria statistica delle classi e calcolo delle probabilita, Pubblicazioni Del R Istituto Superiore Di Scienze Economiche e Commericiali Di Firenze. 8 (1936) 3–62.

[21] M. Allen, D. Poggiali, K. Whitaker, T.R. Marshall, J. van Langen, R.A. Kievit, Raincloud plots: a multi-platform tool for robust data visualization [version 2; peer review: 2 approved], Wellcome Open Res. 4 (2021). https://doi.org/10.12688/wellcomeopenres.15191.2.

[22] M. Arvin, M. Mazaheri, M.J.M. Hoozemans, M. Pijnappels, B.J. Burger, S.M.P. Verschueren, J.H. van Dieën, Effects of narrow base gait on mediolateral balance control in young and older adults, Journal of Biomechanics. 49 (2016) 1264–1267. https://doi.org/10.1016/j.jbiomech.2016.03.011.

[23] T. Ijmker, C.J.C. Lamoth, H. Houdijk, M. Tolsma, L.H.V. van der Woude, A. Daffertshofer, P.J. Beek, Effects of handrail hold and light touch on energetics, step parameters, and neuromuscular activity during walking after stroke, Journal of NeuroEngineering and Rehabilitation. 12 (2015) 70. https://doi.org/10.1186/s12984-015-0051-3.

[24] T. Ijmker, S. Noten, C.J. Lamoth, P.J. Beek, L.H.V. van der Woude, H. Houdijk, Can external lateral stabilization reduce the energy cost of walking in persons with a lower limb amputation?, Gait & Posture. 40 (2014) 616–621. https://doi.org/10.1016/j.gaitpost.2014.07.013.

[25] J.M. Donelan, D.W. Shipman, R. Kram, A.D. Kuo, Mechanical and metabolic requirements for active lateral stabilization in human walking, Journal of Biomechanics. 37 (2004) 827–835. https://doi.org/10.1016/j.jbiomech.2003.06.002.

[26] Y. Wang, M. Srinivasan, Stepping in the direction of the fall: the next foot placement can be predicted from current upper body state in steady-state walking., Biol. Lett. 10 (2014) 20140405. https://doi.org/10.1098/rsbl.2014.0405.

[27] M. Vlutters, E.H.F. van Asseldonk, H. van der Kooij, Center of mass velocity-based predictions in balance recovery following pelvis perturbations during human walking, J. Exp. Biol. 219 (2016) 1514–1523. https://doi.org/10.1242/jeb.129338.

[28] A.L. Hof, J. Duysens, Responses of human hip abductor muscles to lateral balance perturbations during walking, Exp. Brain Res. 230 (2013) 301–310. https://doi.org/10.1007/s00221-013-3655-5.

[29] T.D.J. Welch, L.H. Ting, A feedback model reproduces muscle activity during human postural responses to support-surface translations, J Neurophysiol. 99 (2008) 1032–1038. https://doi.org/10.1152/jn.01110.2007.

[30] S.B. Swart, R. den Otter, C.J.C. Lamoth, Anticipatory control of human gait following simulated slip exposure, Sci Rep. 10 (2020) 9599. https://doi.org/10.1038/s41598-020-66305-1.

[31] T.J.W. Buurke, C.J.C. Lamoth, L.H.V. van der Woude, A.L. Hof, R. den Otter, Bilateral temporal control determines mediolateral margins of stability in symmetric and asymmetric human walking, Sci. Rep. 9 (2019) 12494. https://doi.org/10.1038/s41598-019-49033-z.

[32] J.M. Donelan, R. Kram, A.D. Kuo, Mechanical and metabolic determinants of the preferred step width in human walking, Proc. R. Soc. B. 268 (2001) 1985–1992. https://doi.org/10.1098/rspb.2001.1761.

[33] M. Ferrarin, M. Rizzone, L. Lopiano, M. Recalcati, A. Pedotti, Effects of subthalamic nucleus stimulation and l-dopa in trunk kinematics of patients with Parkinson’s disease, Gait & Posture. 19 (2004) 164–171. https://doi.org/10.1016/S0966-6362(03)00058-4.

[34] D.S. Peterson, M. Mancini, P.C. Fino, F. Horak, K. Smulders, Speeding Up Gait in Parkinson’s Disease, Journal of Parkinson’s Disease. 10 (2020) 245–253. https://doi.org/10.3233/JPD-191682.

[35] H. Xu, M. Hunt, K. Bo Foreman, J. Zhao, A. Merryweather, Gait alterations on irregular surface in people with Parkinson’s disease, Clinical Biomechanics. 57 (2018) 93–98. https://doi.org/10.1016/j.clinbiomech.2018.06.013.

[36] D. De Bartolo, G. Morone, G. Giordani, G. Antonucci, V. Russo, A. Fusco, F. Marinozzi, F. Bini, G.F. Spitoni, S. Paolucci, M. Iosa, Effect of different music genres on gait patterns in Parkinson’s disease, Neurol Sci. 41 (2020) 575–582. https://doi.org/10.1007/s10072-019-04127-4.

[37] J.L. McKay, K.C. Lang, S.M. Bong, M.E. Hackney, S.A. Factor, L.H. Ting, Abnormal center of mass feedback responses during balance: A potential biomarker of falls in Parkinson’s disease, PLOS ONE. 16 (2021) e0252119. https://doi.org/10.1371/journal.pone.0252119.

